# Landscape scale ecology of *Tetracladium spp*. fungal root endophytes

**DOI:** 10.1101/2022.05.26.493577

**Authors:** Anna Lazar, Ryan M. Mushinski, Gary D. Bending

## Abstract

**Background:** The genus *Tetracladium* has been traditionally regarded as an Ingoldian fungus or aquatic hyphomycete – a group of phylogenetically diverse, polyphyletic fungi which grow on decaying leaves and plant litter in streams. Recent sequencing evidence has shown that *Tetracladium* spp. may also exist as root endophytes in terrestrial environments, and furthermore may have beneficial effects on the health and growth of their host. However, the diversity of *Tetracladium* spp. communities in terrestrial systems and the factors which shape their distribution are largely unknown.

**Results:** Using a fungal community internal transcribed spacer amplicon dataset from 37 UK *Brassica napus* fields we found that soils contained diverse *Tetracladium* spp., most of which represent previously uncharacterised clades. The two most abundant OTUs, related to previously described aquatic *T. furcatum* and *T. maxilliforme*, were enriched in roots relative to bulk and rhizosphere soil. For both taxa, relative abundance in roots, but not rhizosphere or bulk soil was correlated with *B. napus* yield. The relative abundance of *T. furcatum* and *T. maxilliforme* OTUs across compartments showed very similar responses with respect to agricultural management practices and soil characteristics. The factors shaping the relative abundance of *T. furcatum* and *T. maxilliforme* OTUs in roots was assessed using linear regression and structural equation modelling. Relative abundance of *Tetracladium maxilliforme* and *Tetracladium furcatum* in roots increased with pH, concentrations of phosphorus, and increased rotation frequency of OSR. While it decreased with increased soil water content, concentrations of extractable phosphorus, chromium, and iron.

**Conclusions:** The genus *Tetracladium* as a root colonising endophyte is a diverse and wildly distributed part of the oilseed rape microbiome that positively correlates to crop yield. The main drivers of its community composition are crop management practices and soil nutrients.

## Background

Aquatic hyphomycetes or Ingoldian fungi are important decomposers in freshwater ecosystems [1]. Spores of these fungi were first described from running freshwater streams in the 1940s [2], with species classified according to morphology – primarily sigmoid or tetraradiate [3]. Sexual reproduction of these fungi has never been observed, and the members of this group don’t share common morphological or ecological characteristics [4]. However, the common conidial shape suggests convergent evolution, and may contribute to spore dispersal via improved anchoring to the substrate or higher buoyancy for better aquatic dispersal [5]. Recently, use of next-generation sequencing has revealed the presence of aquatic hyphomycetes in fungal communities inhabiting soil and plants, although the ecological importance of these fungi in terrestrial habitats is unknown [6].

The genus *Tetracladium* is a common aquatic hyphomycete that was first described by de Wildeman in 1893 [7], and sits within the Ascomycete class Leotiomycetes in the Han Clade 9/Stamnaria lineage/Vandijckellaceae clade as *incertae sedis* [8]. The genus name was coined in response to its distinct ~60×100μm tetra formatted conidiospores which have a central axis with three radiating branches [9]. Since the initial description of *Tetracladium*, the genus has been found to be ubiquitous in aquatic environments [9–14]. The first terrestrial observations of *Tetracladium* were from forest litter [11, 15, 16], with fungal spores detected in the water film covering fallen leaves [15]. However, most reports of *Tetracladium* in terrestrial environments came after the turn of the century as DNA sequencing techniques became more easily accessible. Most of this data comes from environmental metabarcoding studies, and there are only a few instances of *Tetracladium* ssp. being isolated in pure cultures. It has been hypothesised that there may have been under-reporting of *Tetracladium* in terrestrial habitats before the 2000s because of the strange nature of finding an aquatic organism in a terrestrial environment [6]. It is unclear whether the species described based on spore morphology from aquatic habitats and the DNA sequences identified from terrestrial environmental samples belong the same organisms. However ITS (internal transcribed spacer) amplicon analysis has shown no sequence-based differences between aquatic and terrestrial strains of a number of species, indicating that some species may have diverse ecological functions [6].

One of the first observations of plant endophytic *Tetracladium* sp. came from riparian plant roots [17]. These fungi don’t appear to show host or habitat specificity as plant endophytes, and have been found in roots of monocot species within Asparagales [18–20], Liliales [21] and Poales [22–24], and dicot species within Ericales [25], Brassicales [26, 27] and Vitales [28]. Furthermore they have been found associated with Equisetaceae [29, 30] and Bryophytes [31–34]. *Tetracladium* has most frequently been described in metabarcoding studies of soil from disturbed agricultural and grassland habitats [35–38]. There are numerous reports of the genus from the Antarctic tundra [39–42] where they have also been found in unvegetated habitats including glaciers and bedrock [43–45].

The dual ecology of *Tetracladium* sp., and particularly their importance as plant endophytes, are still debated. Anderson et al. [9] investigated an aquatic *T*. *marchalianum* population over time and space and found that the fungus maintains a high genotypic diversity throughout the year, and suggested that this could be attributed to their alternative lifestyles as terrestrial plant endophytes [9]. It was proposed by Selosse et al. (2008) that the terrestrial occurrence of aquatic hyphomycetes, and more specifically their endophytic nature, is attributed to the fungi precolonising plant tissues and building biomass, so in the event of abscission, they are already occupying the niche, and ready to decompose plant litter which reaches freshwater. It was also suggested that the tetraradiate spore morphology could aid them in becoming airborne [6]. Based on this theory *Tetracladium* sp. should be most common in aerial plant tissues, however there is currently no evidence to suggest this is true.

There is conflicting evidence on whether *Tetracladium* infection provides benefits to the host. Glasshouse experiments have shown that inoculation with *Tetracladium* sp. can have beneficial effects on plant growth [46], while other studies have shown no effects [47]. Importantly, Hilton et al. (2021) found that *Tetracladium* sp. had a co-exclusion relationship with root pathogenic fungi, and relative abundace in roots was positively associated with crop yield [38]. Furthermore, *T. marchalianum* showed an antagonistic effect against bacterial plant pathogens including *Erwinia chrysanthemi and Xanthomonas phaseoli*, although other *Tetracladium sp*. showed no such effects [48].

To date there have been no systematic studies which have investigated the diversity and distribution of *Tetracladium* in terrestrial habitats, and as a result the factors which shape *Tetracladium* spp. communities, and the extent to which they interact with plants are unclear. In the current study we build on our earlier work [38] which characterised root fungal communities of *Brassica napus* across 37 UK fields to investigate (1) the diversity of *Tetracladium* spp. in soil and roots at the landscape scale (2) the extent to which different *Tetracladium* spp. are selectively recruited into roots and rhizosphere soil from bulk soil (3) the relationships between root, rhizosphere and soil populations of different *Tetracladium* spp. and crop yield and (4) to determine the importance of, and interactions between, soil nutrients, climate, soil physical properties, and crop management practices as drivers for the colonisation of roots by *Tetracladium* spp.

## Methods

### Sample collection and analyses

Root, rhizosphere soil and bulk soil samples were collected in March 2015 from 37 oilseed rape *(B. napus)* fields from 25 commercial farms in the UK. Following DNA extraction, the fungal community was amplified using internal transcribed spacer primers [49]. Sequencing was performed with Illumina MiSeq technology, and taxonomy assigned using Quantitative Insights into Microbial Ecology (QIIME 1.8) [50] with the UNITE ITS database [51]. Sequences were clustered to OTU at 97 % minimum identity threshold, and those OTUs assigned as *Tetracladium* spp. were selected for use in the current study. Metadata collected from each field included soil physico-chemical parameters (including C, N, P, micronutrient, pH, and soil type), climatic data, crop variety, rotation sequence and grain yield at the subsequent harvest. Full methodological details for sample preparation, DNA extraction, sequencing and bioinformatic analysis can be found in Hilton et al. (2021). Information about farm locations, *Tetracladium* spp. OTUs and metadata can be found in Supplementary Table 1.

### Phylogenetic analyses

To obtain more detailed information about the phylogenetic relatedness of recovered *Tetracladium* spp. sequences, the most closely related sequences to these OTUs were downloaded, including two representative ITS sequences from all described species. Sequences were accessed from the NCBI GenBank, and were aligned with our *Tetracladium* OTU sequences using the MAFFT v.7 (e-ins-I algorithm) [52]. Maximum likelihood analyses were performed with RAxML on the CIPRES Science Gateway to build a phylogenetic tree using the default setting with 1000 bootstraps [53, 54].

### Statistical analyses

Observed species counts were used to generate richness plots to study OTU abundance differences in the three sampled compartments (bulk soil, rhizosphere, and root) using *vegan* in R [55]. Significance of differences in taxa richness between compartments and differences in OTU relative abundance between crop genotypes and previous cultivated crops were tested using the Kruskal–Wallis rank sum test. P values were corrected for multiple comparisons with a Dunn’s test using the false discovery rate with the Benjamini–Hochberg method. Linear regression was used to corelate relative abundance to yield and rotation. Zero values were introduced to accommodate for fields that never had oilseed rape sown before, therefore rotation length values are reciprocal. Ternary plots were created using *ggtern* [56] to understand the compartment preference of the OTUs.

Out of all the OTUs found that resembled the genus *Tetracladium* we chose the two most abundant OTUs, which were also substantively enriched within the root compartment, for detailed ecological analyses. Data was normalised using modified Z-scores. To test for drivers of relative abundance in the roots, we created a piecewise structural equation model (PSEM). First, a correlogram was created to better understand relationships between metadata and OTU relative abundance (**Additional file 1**) using base R functions [57]. Then, individual linear mixed-effect models were fitted with sampling location and soil compartment as random variables. Significant soil nutrient factors, relative to OTU abundance, taken from the correlogram output and the initial fitted model with all soil nutrients (modM) (**Additional files 1 and 2**), soil structure and climate were used as composite fixed variables. These composite variables were created so complicated constructs can be processed as simpler blocks that are easier to present and discuss [58]. Variable reduction was done to unmask significant relationships that may be missed if too many factors were included based on the LMM fitted with all variables. Individual models were fitted using the R package *lme4* [59]. Fixed variables were reduced via assessing best model fit using the *performance* package [60]. Finally, path models were fitted with the *piecewiseSEM* package [61], based on the findings of the individual models.

## Results

Across the 37 fields we found twelve OTUs that represented the genus *Tetracladium* (**Fig. 1A**). Higher abundance *Tetracladium* sp. OTUs (OTUs 19, 1088, 168, 6663, 359, 4156, 5882) were found in all sampled fields, while lower abundance OTUs (813, 1055, 3952, 6656, 10312) were found sporadically across fields (**Additional file 3C**). The highest abundance OTUs, OTU 19 and 1088, grouped with *T*. *maxilliforme* and *T. furcatum* respectively, both of which have been described from water (**Fig. 1B**). OTU 813 clustered closely with, but was not identical to the species *T. elipsoideum*, which has been described from arctic soil [44] and the closest uncultured environmental sequences to this OTU (MF181805.1 and MK627297.1) also originate from soil. OTUs 6663, 5882, 4156, and 168 clustered with uncultured *Tetracladium* sp. sequences in a clade close to *T.ellipsoideum, T. psychrophilum* and *T. globosum*. The closet uncultured environmental sequences for OTU 6663 (LR863329.1 and KX192428.1), OTU 5882 (MN660389.1), OTU 4156 (KF296960.1 and JX029127.1), OTU 168 (GU055746.1) all originate from terrestrial samples. Environmental sequence MN660932.1 found in water, was a close match to OTU 168. OTUs 1055, 3952, 6656, 10312, and 359 formed a distinct clade with uncultured *Tetracladium* sp. sequences (MW050202.1, MH451254.1, KX193670.1, MH451294.1, MG756632.1, KM246272.1, MK246209.1, LR876862.1, MK627349.1) found on land, likely representing uncharacterised species.

**Figure 1.**
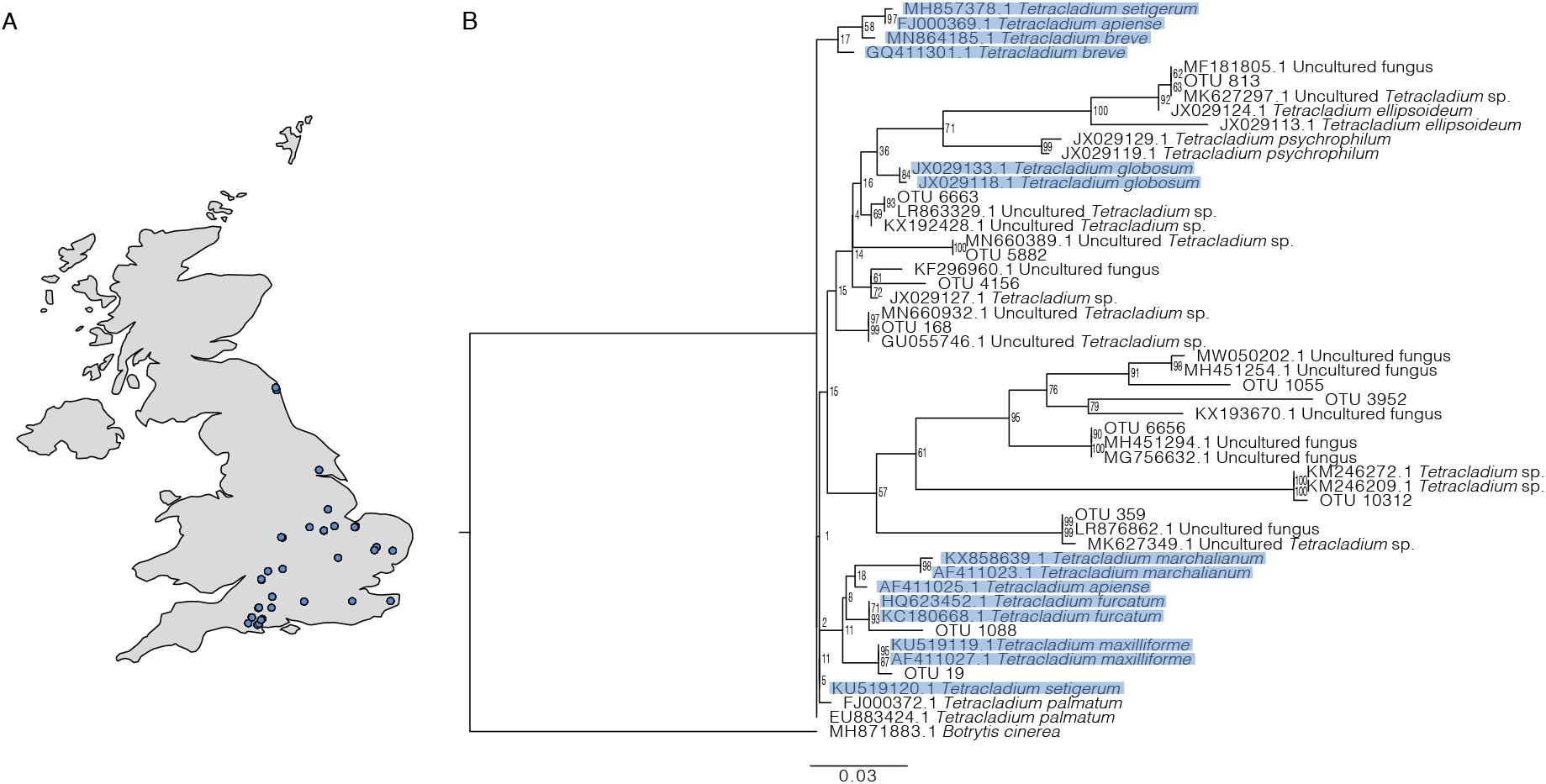
A - Location of the sampling sites used in this study. B – ITS sequence based maximum likelihood tree with posterior probability values of the twelve *Tetracladium* sp. OTUs and reference sequences. *Botrytis cinerea* was used as an outgroup. The scale bar denotes the number of nucleotide differences per site.

There was a significant difference between observed OTU richness of the three compartments (**Fig. 2A**), which was the lowest in the roots while the bulk soil and rhizosphere showed the same OTU richness (rhizosphere-bulk soil *P* = <0.001, root-bulk soil *P* = <0.001, root-rhizosphere *P* = <0.001). Five of the OTUs (OTUs 19, 5882, 1055, 813, 1088) were most abundant in the roots, while OTU 6656 was found in greater abundance in the bulk soil, and OTU 10312 was only found in the rhizosphere. The rest of the OTUs didn’t show a specific preference for compartment (**Fig. 2B**). The most abundant *Tetracladium* OTUs (19 and 1088) both showed a strong preference for the roots. The mean relative abundance of OTU 19 was five times higher in the roots than in the bulk soil and it was over a thousand times higher in the case of OTU 1088 (**Additional file 3D**).

**Figure 2.**
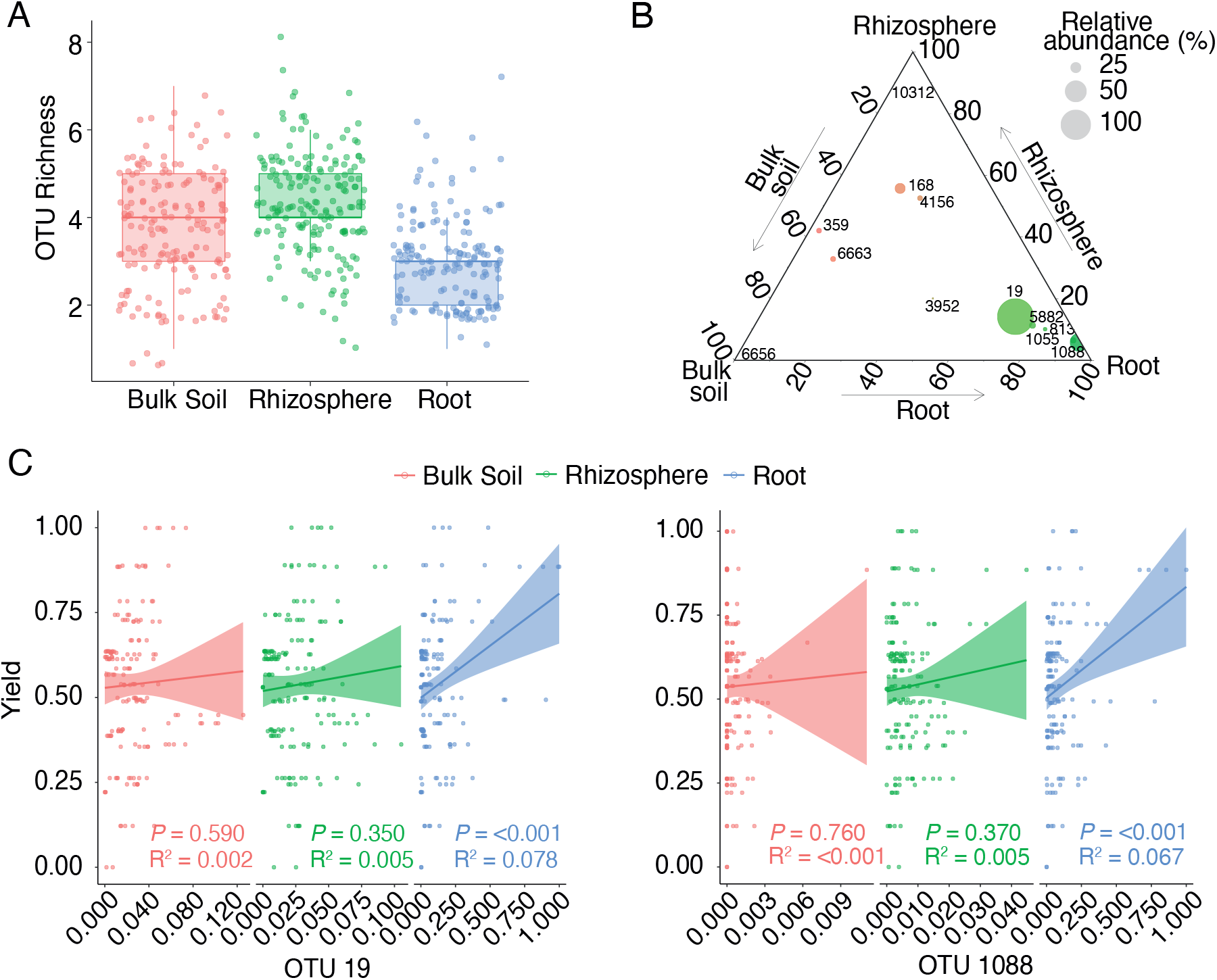
Analyses of *Tetracladium* OTU distribution in the bulk soil, the rhizosphere, and the roots. A – Observed OTU richness in the three sampled compartments. Error bars represent standard error. B – Ternary plots of *Tetracladium* OTU distribution across compartments. C – Linear regression line fitted between OTUs and yield across the three compartments with significance values. OTU relative abundances and yield are normalised. The shaded region represents the 95% confidence limits for the estimated prediction.

Based on these results we conclude that OTUs 19 and 1088 can be categorised as root colonising fungi and further analyses focussed on these two abundant and widely distributed OTUs.

### Correlating the metadata to OTU relative abundance

Relative abundance of OTUs 19 and 1088 in the root compartment had a significant positive linear correlation with OSR yield (OTU 19 – R^2^ = 0.078, *P* = <0.001, OTU 1088 – R^2^ = 0.067, *P* = <0.001). The highest relative abundance of OTU 19 and 1088 in the roots was associated with a yield increase of up to 25% relative to samples with the lowest relative abundance. No such relationship was seen in the bulk soil or rhizosphere (**Fig. 2C**).

We further investigated the relationships between previous crop cultivated at the site, OSR variety and rotation and the relative abundance of OTUs 19 and 1088. We found that relative abundance of OTU 19 and 1088 had a significant correlation with previous cultivated crop (R^2^ = 0.021 *P* = 0.013 and R^2^ = 0.021, *P* = 0.012 respectively). In roots, relative abundance of both OTU 19 and 1088, was significantly higher (*P* <0.001) when rye rather than barley or wheat was the proceeding crop **(Fig. 3A)**. In the rhizosphere and soil relative abundance of OTU 19 showed the same patterns in relation to previous crops as for the roots, but in addition relative abundance following fallow was significantly higher than barley and lower than wheat (**Figs. 3A** and **3B**, **Additional file 4**).

**Figure 3.**
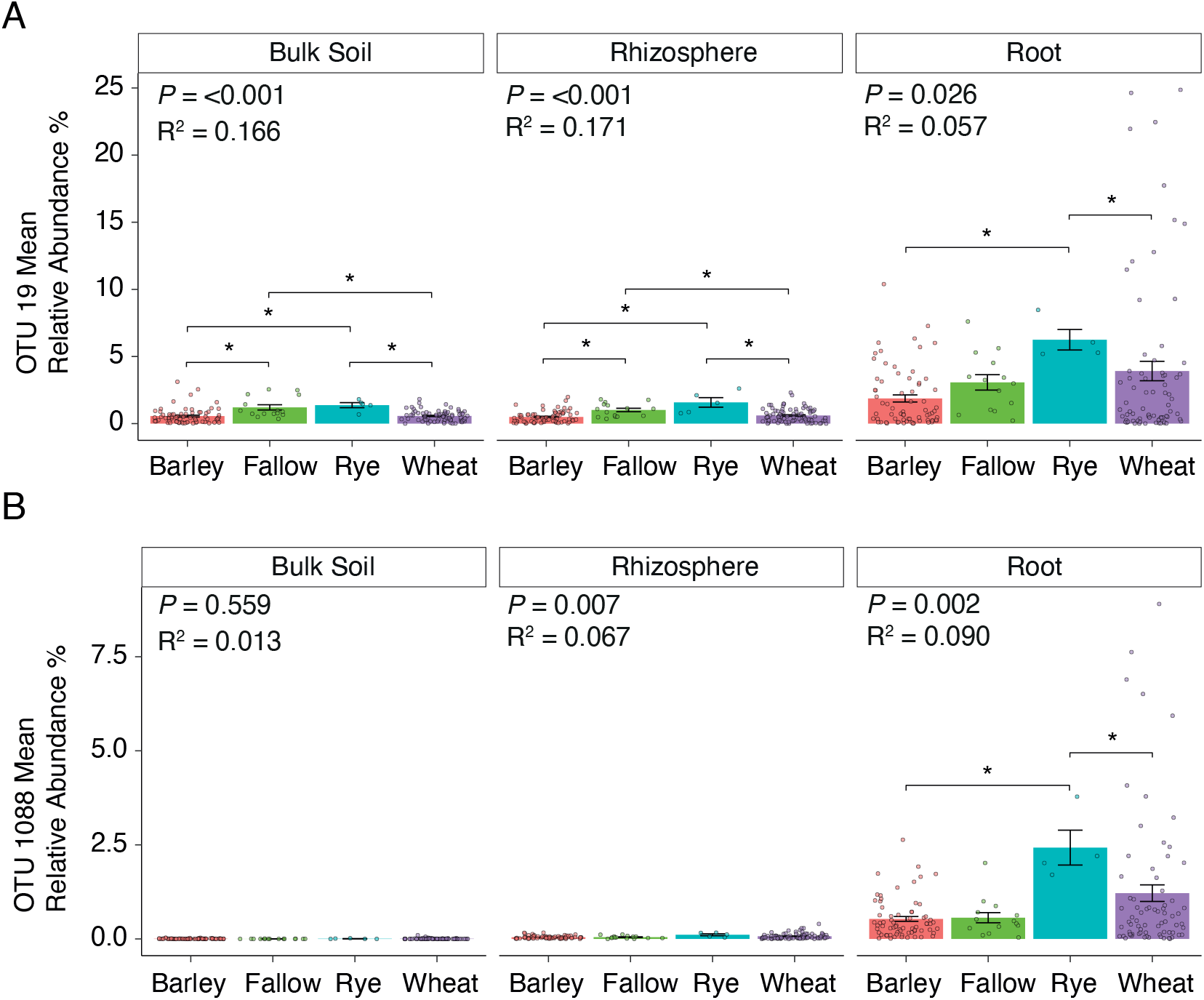
The mean relative abundance of A – OTU 19 and B – OTU 1088 in the four previous crop types in the bulk soil, the rhizosphere soil, and the roots. Stars indicate a significant difference between the previous crops. Error bars represent +/− standard error of the mean.

OSR variety had a significant effect on relative abundance of OTU 19 (R^2^ = 0.615, *P* = <0.001) and 1088 (R^2^ = 0.614, *P* = <0.001) in roots, with the two OTUs showing very similar distribution patterns. Seven varieties (Quartz, Rocca, Incentive, Compass, Catena, Camelot and Cabernet) had very low mean relative abundances of these OTUs, while Nikita had substantively higher relative abundance than the other varieties (**Figs. 4A** and **4B**, **Additional file 5**). Rotation also had a significant linear correlation with the relative abundance of OTUs 19 and 1088 in the roots (R^2^ = 0.032, *P* = 0.023 for OTU 19, R^2^ = 0.059, *P* = 0.002 for OTU 1088) (**Fig. 4B**), which increased as time since previous OSR crop increased.

**Figure 4.**
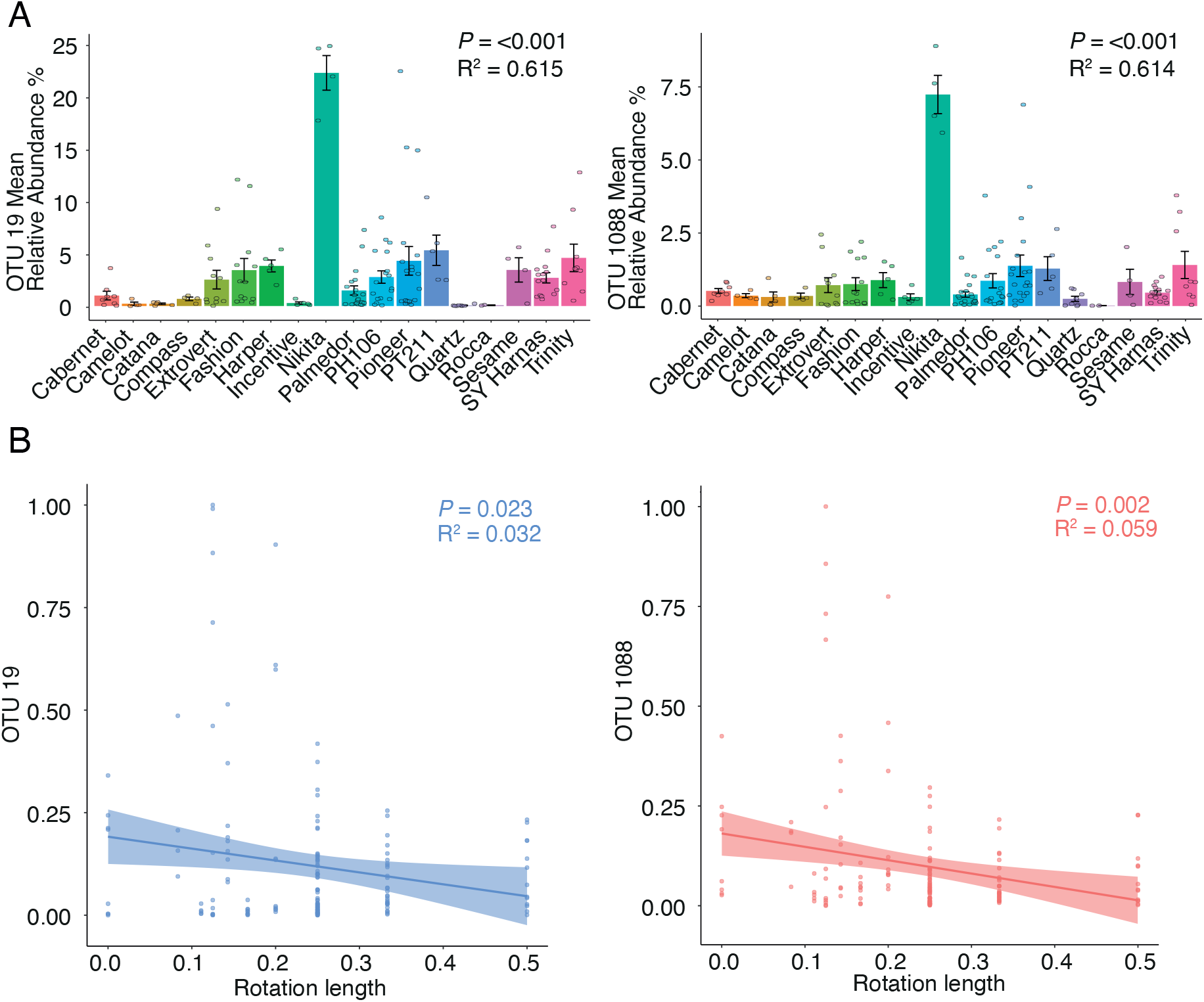
Relationships between *Tetracladium* spp. OTU mean relative abundance with variety and rotation. A - The mean relative abundance of OTU 19 and 1088 in different OSR varieties in the roots. Error bars represent +/− standard error of the mean. B – Linear regression line fitted between OTU relative abundance and rotation length across the three compartments with significance values. OTU relative abundances and yield are normalised in a way to accommodate for fields that never had oilseed rape planted before. These virgin fields are represented as 0 while the shortest rotation length is represented by 0.5. The shaded region represents the 95% confidence limits for the estimated prediction.

We found a strong correlation between the relative abundances of the root specific *Tetracladium* OTUs (R^2^ = 0.887, *P* = <0.001) so for further analyses we used the mean of the combined relative abundance of the two OTUs of each sample (**Fig. 5**). In the final PSEM, we correlated relative abundance of the combined OTUs to soil structure (soil moisture content, bulk density, and sand content), significant nutrients (Olsen P, iron, chromium, phosphorus, and manganese), pH, rotation, and climate (annual rainfall and minimum annual temperature) using sampled field as a random variable and used simple linear regression to correlate nutrients to soil structure and then pH to nutrients. Nutrient variables were chosen based on the results of the correlation matrix and the assessment the model fit of the initial linear mixed-effect model modM (**Additional file 1**).

**Figure 5.**
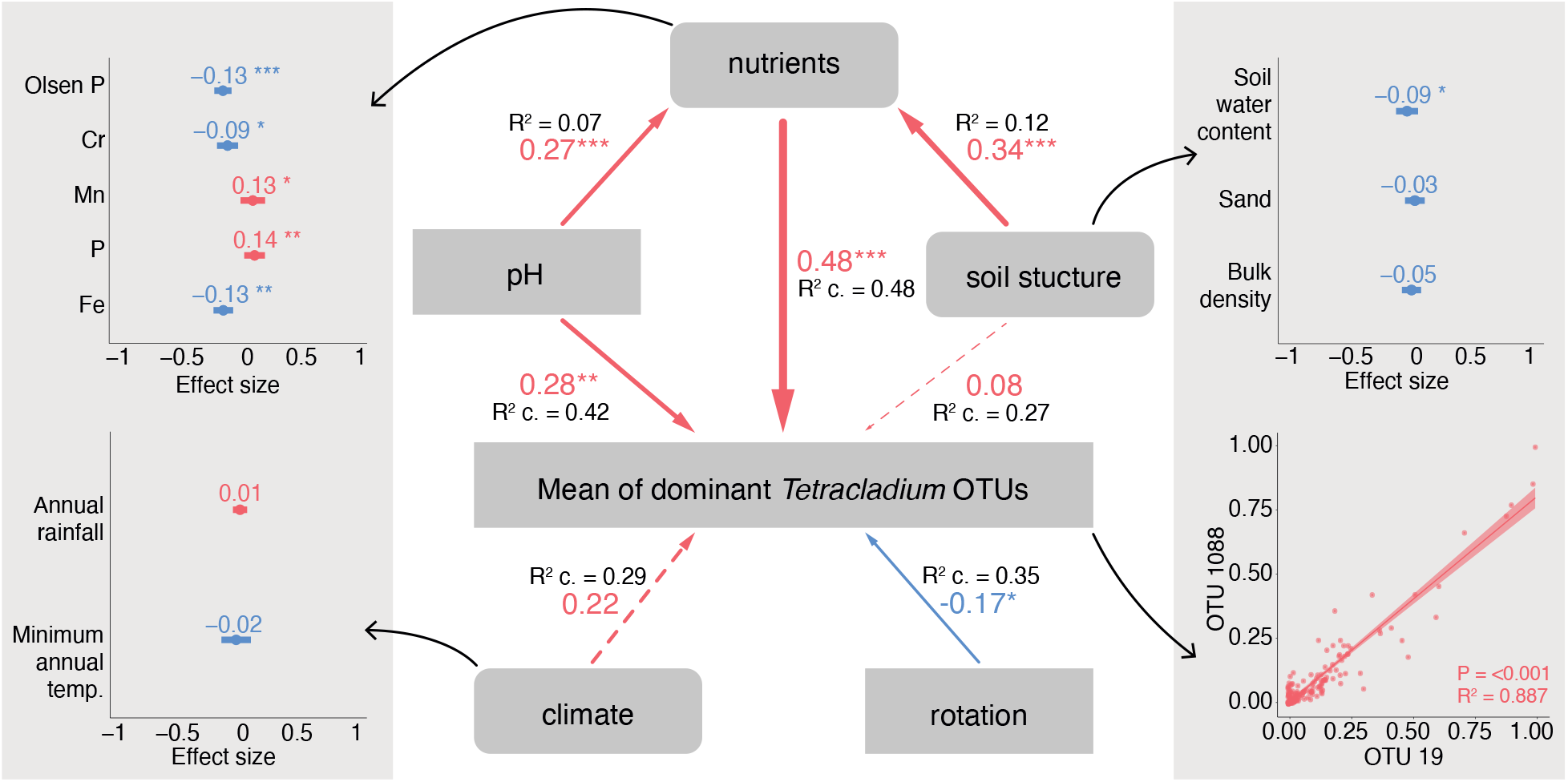
Drivers of the combined mean relative abundance of the two OTUs. Path diagrams of the piecewise SEMs showing direct and indirect effects with standard estimates R^2^ values for linear correlations and R^2^ conditional values for mixed effects linear correlations. The R2 marginal values for nutrients, pH, soil structure, climate, rotation are: 0.35, 0.20, 0.03, 0.01, 0.03. Dotted lines indicate non-significance in the path model. Arrow sizes indicate effect size, arrow colours indicate a positive or a negative relationship (red – positive, blue – negative). The multivariate PSEM linking soil structure, pH, rotation, climate, and soil nutrients with mean relative abundance of OTUs 19 and 1088 was well supported by the data (Fisher’s C = 36.86, *P* = <0.001, degrees of freedom = 10). *p < 0.05, **p < 0.01, ***p < 0.001. Standardised effect size is shown from the linear mixed effects models where soil compartment and sampled location were used as random effect. Model fit indicators for the LMMs are shown in Additional file 2. Rotation values are reciprocal.

As a result, the PSEM showed the strongest significant direct positive effect of nutrients (R^2^ conditional = 0.48, standard estimate = 0.48, *P* = 0.001) followed by pH (R^2^ conditional = 0.07, standard estimate = 0.27, *P* = 0.002) and rotation (R^2^ conditional = 0.35, standard estimate = 0.17, *P* = 0.043) on the mean combined relative abundance of OTU 19 and 1088 (**Fig. 5**). The effect of climate and soil structure was not significant in the path analyses. Soil structure and pH had significant positive correlations with nutrients (R^2^ = 0.12, standard estimate = 0.34, *P* =< 0.001 and R^2^ conditional = 0.07, standard estimate = 0.27, *P* = 0.005) thus indirectly affecting OTU relative abundance. To expand the composite variables of the PSEM, the mixed-effect linear models showed significant correlation between the combined mean relative abundance of OTU 19 and 1088 (**Fig. 5**) with Olsen P (*P* =< 0.001), phosphorus (*P* = 0.003) and iron (*P* = 0.008). We also found a significant negative correlation between the combined mean relative abundance of the OTUs and soil water content (*P* = 0.045), while none of the climate variables were significant (**Fig. 5**).

## Discussion

Here we present the first systematic study of the landscape scale diversity and distribution of *Tetracladium* spp. within terrestrial systems and identify key factors controlling their occurrence as root endophytes. *Tetracladium* spp. were widely distributed, occurring in soil, rhizosphere, and roots in the 37 sampled sites. A total of 12 *Tetracladium* sp. OTUs were detected, and a subset of OTU were specifically enriched in oilseed rape roots, including *T. maxilliforme* and *T. furcatum*. Eight of the OTUs belonged to clades for which only environment sequences, largely from terrestrial habitats, have been described. There was a significant relationship between relative abundance of *T. maxilliforme* and *T. furcatum* within oilseed rape roots and crop genotype, previous cultivated crop, and oilseed rape rotation period. Linear mixed effects modelling, and piecewise structural equation modelling showed that the most important environmental drivers of the relative abundance of *Tetracladium* spp. within plant roots were pH and select nutrients, including total phosphorus, extractable (Olsen) phosphorus, and iron.

### Diversity and distribution

*Tetracladium* is generally considered to be a freshwater fungus, however, it has also been detected as a root endophyte of terrestrial plants. In the sampled fields there was lower species richness in the roots, than in the soil and rhizosphere which is a common feature of endophytes, and suggests selective recruitment into the microbiome [62]. Based on ITS sequences, the two most abundant OTUs (OTU 19 and 1088) clustered with *T. maxilliforme* and *T. furcatum* respectively and showed a strong root preference (**Fig. 2B**). *T.maxilliforme* and *T. furcatum* have been found several times in agriculture including the roots of crop plants [22, 25], though they are both traditionally regarded as aquatic organisms [14, 63–67]. OTU 813 clustered with the terrestrial *T. elipsoideum* and it showed a strong root preference (**Fig. 2B**). Excluding OTUs 19, 1088, and 813, the closest GenBank sequence matches for all OTUs were from terrestrial habitats. Compartment preference did not have a correlation with taxonomic position as root preferring OTUs were found across the main *Tetracladium* spp. clades suggesting that the different species in the genus may not have a defined lifestyle. Overall, our findings suggest that there is considerable diversity within *Tetracladium* spp. and that while some taxa may inhabit both aquatic and terrestrial habitats, others may inhabit terrestrial systems, with some occurring as plant endophytes.

Root endophytes have been shown to increase host resistance to environmental factors such as drought, heat and saline stress [68, 69], and can also increase plant health by inducing increased resistance via priming of the natural immune system to pathogens [70] or through competition with pathogens for nutrients and spatial niche exclusion [71]. Plant growth and yield can also be increased by endophytes via direct nutrient transfer from fungus to plant [72]. Furthermore, root colonising endophytes may shape plant community diversity and distribution [73].

Although, *Tetracladium* species have previously been found in roots [19, 22, 25, 26] little is known about the process by which *Tetracladium* spp. colonise roots, or the significance of infection for plant health. Sati and Arya (2010) found that inoculation with *T. nainitalense* had no significant effect on the growth of *Hibiscus esculentus* or *Solanum melongena* following inoculation with *T. nainitalense*, although there was no evidence that the inoculant colonised plant tissues [47]. In our previous work [38] we found a positive correlation between *Tetracladium* OTU relative abundance and oilseed rape yield on a landscape scale and in the current study we build on this to show a 25% yield increase from the lowest to the highest OTU relative abundance with both OTUs 19 and 1088. There is a clear need to understand the root infection process by *Tetracladium* spp. and to quantify benefits for plant health under controlled conditions, so that the significance of *Tetracladium* spp. root endophytes, and their potential to act as beneficial symbionts can be established.

### Drivers of relative abundance

Here we present the first study that systematically investigates the drivers of root colonisation by *Tetracladium* spp. The root associated *T. maxilliforme* and *T. furcatum* OTUs showed strong co-assembly patterns in oilseed rape roots at a landscape scale. These OTUs showed the same interactions with host genotype, crop management, and environmental factors. There is evidence of minimal competition between root colonising endophytes explaining their high diversity within a single host [74]. This enables similar strains or species of fungi to colonise the same plant in high abundance. The data presented here could simply mean similar adaptation of the two species without any ecological interaction.

According to our fitted models, the main drivers of *Tetracladium* sp. relative abundance in oilseed rape roots were soil nutrient content, crop rotation and pH. Relative abundance had a positive correlation with soil phosphorus and a negative correlation with iron content and Olsen P. Phosphorus and iron availability may limit plant and microbial growth in soil. For example, dark septate endophytes, which like *Tetracladium* spp. belong to the Helotiales, and occur as root endophytes, have been found to have iron phosphate solubilisation properties [75]. Sati and Pant found phosphate solubilisation in *T. setigerum* isolated from riparian roots in agar and broth media [76]. Moreover, it was shown that mineral fertilisers increased the relative abundance of *Tetracladium* sp. indicating that mineral fertiliser treatment might promote this plant-fungal symbiosis [23, 77]. In our study, total phosphorus content of the soil had a positive relationship with *Tetacladium* spp. OTU relative abundance in roots. In contrast, extractable phosphorus or Olsen P had a negative relationship with *Tetracladium* spp. OTU relative abundance in roots. This could suggest that *Tetracladium* spp. infection is promoted when bioavailability of P is low, in the same way that plants favour colonisation by arbuscular mycorrhizal symbioses under conditions of low P availability [78].

In contrast to bacteria, fungal communities are favoured by low soil pH [79]. However, *T. maxilliforme* and *T. furcatum* OTUs showed increased relative abundance in roots as soil pH increased. Furthermore, pH was a key factor in determining fungal community structure in the soil in many cases where *Tetracladium* has been identified as a common genus [41, 80], however it was found to prefer neutral or slightly acidic soil in a long-term microplot experiment [80]. In addition to pH, soil redox potential (Eh) is an important driver of microbial community growth, diversity and composition [81, 82]. Relative abundance of both *T. maxilliforme* and *T. furcatum* increased as soil moisture decreased, which was surprising considering their dual ecology as aquatic taxa. Low soil moisture combined with high pH leads to lower Eh in the soil and results in slower rates of decomposition [83, 84]. In addition, high soil moisture and high Eh result in increased reducing conditions that limit extractable P availability in the soil through the solubilisation of Fe oxides that bind to available P [85]. Root colonising *Tetracladium* OTUs showed higher relative abundance in low reducing conditions (low soil moisture and high pH) and had higher relative abundance under low P availability, however this data originates from bulk soil physicochemical measurements and therefore the results may differ when looking at the soil closely encapsulating the roots.

Finally, we found a strong correlation between OTU relative abundance and crop management practices. OTUs 19 and 1088 both had the highest relative abundances in all compartments when OSR was planted after rye and their relative abundance increased with OSR rotation length. Crop rotation is known to influence soil microbial community composition, and to influence the composition of plant associated microbiota including pathogens and symbionts [86, 87]. These changes may be attributed to a wide range of interactions. For endophytes, they may reflect plant species specificity, and the extent to which different crop species support proliferation of inoculum, either following recruitment in living root biomass or on organic material left in the field following harvest [88]. Additionally, differences in management practices across crop types, such as fertiliser, tillage and pesticide use could also impact inoculum (Gosling et al., 2006). Enhanced colonisation following ryegrass relative to OSR could therefore indicate a preference of *Tetracladium* spp. for ryegrass as a host, or management practices assisted with ryegrass.

## Conclusion

The ecological interactions of *Tetracladium spp*. are a currently unknown; however, there is overwhelming evidence that some taxa within this group which were traditionally considered to be aquatic hyphomycetes can also occur as endophytes in terrestrial ecosystems, with several clades known only from environmental DNA, and which may represent terrestrial species. We have found a correlation between crop yield and *Tetracladium* abundance, indicating that these fungi are beneficial components of the plant endophytic mycobiome. There is also indication that crop management practices, pH and nutrient enrichment are the main drivers of root colonisation of *Tetracladium* sp. in terrestrial environments. Further research is needed to determine their role in the plant’s life, particularly their effects on plant health and nutrition, to establish their potential value for utilisation in sustainable agricultural practices.

## Declarations

### Ethics approval and consent to participate

Not applicable

### Consent for publication

Not applicable

### Availability of data and material

All data generated or analysed during this study are included in this published article [and its supplementary information files] and in Hilton, S., Picot, E., Schreiter, S. et al. Identification of microbial signatures linked to oilseed rape yield decline at the landscape scale. Microbiome 9, 19 (2021). https://doi.org/10.1186/s40168-020-00972-0

### Authors’ contributions

GDB and RMM were responsible for project conception, project funding, and experimental design. AL, GDB, RMM analysed and interpreted the data. AL wrote the initial manuscript. GDB, RMM reviewed and edited the manuscript.

## Acknowledgements

Not applicable

## Additional files

Additional file 1. Correlogram showing Pearson’s correlation of all metadata variables and OTU relative abundance. Boxes are coloured according to the R values blue indicating a negative, red indicating a positive relationship. Non-significant relations are shown with an x in the box.

Additional file 2. Model indicators for LMMs. A is a visual representation of the measured model fit indices for the nutrient models. The final model included in figure 5 is modMr. B is the actual values corresponding to the indicators of the nutrient models. C is the model fit indicators of the soil structure and climate models from Figure 5.

Additional file 3. Metadata tables. A - Metadata table from the 25 farms. B – Bulk soil properties from each five reps of the 37 field sites. C - Tetracladium OTU relative abundances across samples. D – Mean Tetracladium OTU relative abundances across the three sampled compartments.

Additional file 4. Dunn’s test results correcting for P values with Benjamini– Hochberg method. Comparing previous crop types across the different compartments in OTU 19 and OTU 1088.

Additional file 5. Dunn’s test results correcting for P values with Benjamini – Hochberg method. Comparing OTU 19 and OTU 1088 relative abundance in different oilseed rape varieties in the roots.

## References

1. Grossart, H.-P., et al., Fungi in aquatic ecosystems. Nature Reviews Microbiology, 2019. 17: p.339–354.

2. Ingold, C.T., Aquatic hyphomycetes of decaying alder leaves. Transactions of the British Mycological Society, 1942. 25(4): p. 339–417.

3. Ingold, C.T., Aquatic Hyphomycetes from Switzerland. Transactions of the British Mycological Society, 1949. 32(3-4): p. 341–345.

4. Johnston, P.R., et al., A multigene phylogeny toward a new phylogenetic classification of Leotiomycetes. IMA Fungus, 2019. 10: p. 1.

5. Webster, J., Experiments with Spores of Aquatic Hyphomycetes: I. Sedimentation, and Impaction on Smooth Surfaces. Annals of Botany, 1959. 24(4): p. 595–611.

6. Selosse, M.-A., M. Vohník, and E. Chauvet, Out of the rivers: Are Some Aquatic Hyphomycetes Plant endophytes? New Phytologist, 2008. 178(1): p. 3–7.

7. de Wildeman, É., Notes mycologiques IV. Annales de Société Belge de Microscopie, 1893. 17(2): p. 35–40.

8. Hibbett, D.S., et al., A higher-level phylogenetic classification of the Fungi. Mycol Res, 2007. 111(Pt 5): p. 509–47.

9. Anderson, J.L. and C.A. Shearer, Population Genetics of the Aquatic Fungus Tetracladium marchalianum over Space and Time. PLoS ONE, 2011. 6(1): p. e15908.

10. Conway, K.E., The Aquatic Hyphomycetes of Central New York. Mycologia, 1970. 62(3): p. 516–530.

11. Makela, K., Some Aquatic Hyphomycetes on Grasses in Finland. Karstenia, 1973. 13: p. 16–22.

12. Chauvet, E., Aquatic Hyphomycete Distribution in South-Western France. Journal of Biogeography, 1991. 18(6): p. 699.

13. Bärlocher, F., The Ecology of Aquatic Hyphomycetes. Ecological Studies 1992: Springer, Berlin, Heidelberg.

14. Harrington, T.J., Aquatic Hyphomycetes of 21 Rivers in Southern Ireland. Biology and Environment: Proceedings of the Royal Irish Academy, 1997. **97B**(2): p. 139–148.

15. Bandoni, R.J., Terrestrial Occurrence of Some Aquatic Hyphomycetes. Canadian Journal of Botany, 1972. 50(11): p. 2283–2288.

16. Bärlocher, F. and J.J. Oertli, Colonization of Conifer Needles by Aquatic Hyphomycetes. Canadian Journal of Botany, 1978. 56(1): p. 57–62.

17. Sati, S.C. and M. Belwal, Aquatic Hyphomycetes as Endophytes of Riparian Plant Roots. Mycologia, 2005. 97(1): p. 45–49.

18. Abadie, J.-C., et al., Cephalanthera longifolia (Neottieae, Orchidaceae) Is Mixotrophic: a Comparative Study between Green and Nonphotosynthetic Individuals. Canadian Journal of Botany, 2006. 84(9): p. 1462–1477.

19. Stark, C., W. Babik, and W. Durka, Fungi from the Roots of the Common Terrestrial Orchid Gymnadenia conopsea. Mycological Research, 2009. 113(9): p. 952–959.

20. Vendramin, E., et al., Identification of Two Fungal Endophytes Associated with the Endangered Orchid Orchis militaris L. Journal of Microbiology and Biotechnology, 2010. 20(3): p. 630–636.

21. Wang, Y., et al., Niche differentiation in the rhizosphere and endosphere fungal microbiome of wild Paris polyphylla Sm. PeerJ, 2020. 8: p. e8510.

22. Grudzinska-Sterno, M., et al., Fungal communities in organically grown winter wheat affected by plant organ and development stage. European Journal of Plant Pathology, 2016. 146(2): p. 401–417.

23. Chen, Z., et al., Fungal Community Composition Change and Heavy Metal Accumulation in Response to the long-term Application of Anaerobically Digested Slurry in a Paddy Soil. Ecotoxicology and Environmental Safety, 2020. 196: p. 110453.

24. Zhao, Y., et al., Variation of Rhizosphere Microbial Community in Continuous mono-maize Seed Production. Scientific Reports, 2021. **11**(1).

25. Bruzone, M.C., S.B. Fontenla, and M. Vohnik, Is the Prominent Ericoid Mycorrhizal Fungus Rhizoscyphus ericae Absent in the Southern Hemisphere Ericaceae a Case Study on the Diversity of Root Mycobionts in Gaultheria spp. from Northwest Patagonia, Argentina. Mycorrhiza, 2014. 25(1): p. 25–40.

26. Macia-Vicente, J.G., B. Nam, and M. Thines, Root filtering, Rather than Host Identity or age, Determines the Composition of root-associated Fungi and Oomycetes in Three Naturally co-occurring Brassicaceae. Soil Biology and Biochemistry, 2020. 146: p. 107806.

27. Chatterton, S., et al., Bacterial and Fungal communities, but Not Physicochemical properties, of Soil Differ According to Root Rot Status of Pea. Pedobiologia, 2021. 84: p. 150705.

28. Ma, W., et al., Microbial Diversity Analysis of Vineyards in the Xinjiang Region Using high-throughput Sequencing. Journal of the Institute of Brewing, 2018. 124(3): p. 276–283.

29. Sati, S.C., P. Arya, and M. Belwal, Tetracladium nainitalense sp. nov., a Root Endophyte from Kumaun Himalaya, India. Mycologia, 2009. 101(5): p. 692–695.

30. Giesemann, P., et al., Dark septate endophytes and arbuscular mycorrhizal fungi (Paris-morphotype) affect the stable isotope composition of ‘classically’ non-mycorrhizal plants. Functional Ecology, 2020. 34(12): p. 2453–2466.

31. Russell, J. and S. Bulman, The Liverwort Marchantia foliacea Forms a Specialized Symbiosis with Arbuscular Mycorrhizal Fungi in the Genus Glomus. New Phytologist, 2004. 165(2): p. 567–579.

32. Rosa, L.H., et al., Endophytic Fungi Community Associated with the Dicotyledonous Plant Colobanthus Quitensis (Kunth) Bartl. (Caryophyllaceae) in Antarctica. FEMS Microbiology Ecology, 2010. 73: p. no–no.

33. Hirose, D., et al., Diversity and Community Assembly of moss-associated Fungi in ice-free Coastal Outcrops of Continental Antarctica. Fungal Ecology, 2016. 24: p. 94–101.

34. Hirose, D., et al., Abundance, richness, and Succession of Microfungi in Relation to Chemical Changes in Antarctic Moss Profiles. Polar Biology, 2017. 40(12): p. 2457–2468.

35. Klaubauf, S., et al., Molecular diversity of fungal communities in agricultural soils from Lower Austria. Fungal Divers, 2010. 44(1): p. 65–75.

36. Liu, Y., et al., Analysis of Microbial Diversity in Soil under Ginger Cultivation. Scientifica, 2017. 2017: p. 1–4.

37. María, T., et al., Fairy Rings Harbor Distinct Soil Fungal Communities and High Fungal Diversity in a Montane Grassland. Fungal Ecology, 2020. 47: p. 100962.

38. Hilton, S., et al., Identification of microbial signatures linked to oilseed rape yield decline at the landscape scale. Microbiome, 2021. 9(1): p. 19.

39. Bridge, P.D. and K.K. Newsham, Soil Fungal Community Composition at Mars Oasis, a Southern Maritime Antarctic site, Assessed by PCR Amplification and Cloning. Fungal Ecology, 2009. 2(2): p. 66–74.

40. Tsuji, M., et al., Cold Adaptation of Fungi Obtained from Soil and Lake Sediment in the Skarvsnes ice-free area, Antarctica. FEMS Microbiology Letters, 2013. 346(2): p. 121–130.

41. Zhang, T., et al., Soil pH is a Key Determinant of Soil Fungal Community Composition in the Ny-Ålesund Region, Svalbard (High Arctic). Front Microbiol., 2016. **7**(227).

42. Durán, P., et al., Occurrence of Soil Fungi in Antarctic Pristine Environments. Frontiers in Bioengineering and Biotechnology, 2019. **7**.

43. Jiang, N., et al., Characteristic Microbial Communities in the Continuous Permafrost beside the Bitumen in Qinghai-Tibetan Plateau. Environmental Earth Sciences, 2015. 74(2): p. 1343–1352.

44. Wang, M., et al., Psychrophilic fungi from the world’s roof. Persoonia, 2015. 34: p. 100–12.

45. Hu, W., et al., Multiple-trophic patterns of primary succession following retreat of a high-elevation glacier. Ecosphere, 2021. **12**(3).

46. Sati, S.C. and P. Pant, Two Root Endophytic Aquatic Hyphomycetes Campylospora parvula and Tetracladium setigerum as Plant Growth Promoters. Asian Journal of Agricultural Research, 2020. 4(1): p. 28–33.

47. Sati, S.C. and P. Arya, Assessment of root endophytic aquatic hyphomycetous fungi on plant growth. Symbiosis, 2010. 50(3): p. 143–149.

48. Arya, P. and S.C. Sati, Evaluation of endophytic aquatic hyphomycetes for their antagonistic activity against pathogenic bacteria International Research Journal of Microbiology, 2011. 2(9): p. 343–347.

49. Ihrmark, K., et al., New primers to amplify the fungal ITS2 region – evaluation by 454-sequencing of artificial and natural communities. FEMS Microbiology Ecology, 2012. 82(3): p. 666–667.

50. Caporaso, J.G., et al., QIIME allows analysis of high-throughput community sequencing data. Nat Methods, 2010. 7(5): p. 335–336.

51. Kõljalg, U., et al., Towards a unified paradigm for sequence-based identification of fungi. Mol Ecol. 2013. 22(21): p. 5271–5277.

52. Katoh, K., J. Rozewicki, and K.D. Yamada, MAFFT online service: multiple sequence alignment, interactive sequence choice and visualization. Brief Bioinform.. 2019. 20(4): p. 1160–1166.

53. Miller, M.A., W. Pfeiffer, and T. Schwartz, Creating the CIPRES Science Gateway for inference of large phylogenetic trees. Proceedings of the Gateway Computing Environments Workshop (GCE), 2010: p. pp. 1–8.

54. Stamatakis, A., RAxML version 8: a tool for phylogenetic analysis and post-analysis of large phylogenies Bioinformatics, 2014. 30(9): p. 1312–1313.

55. Oksanen, J., et al. vegan: Community Ecology Package. 2018; R package version 2.5-7]. Available from: https://CRAN.R-project.org/package=vegan.

56. Hamilton, N.E. and M. Ferry, ggtern: Ternary Diagrams Using ggplot2. Journal of Statistical Software, 2018. 87: p. 1–17.

57. R Core Team, R: A language and environment for statistical computing. R Foundation for Statistical Computing, Vienna, Austria, 2021.

58. Grace, J.B. and J.E. Keeley, A structural equation model analysis of postfire plant diversity in California shrublands. Ecological Applications, 2006. 16(2): p. 503–514.

59. Bates, D., et al., Fitting Linear Mixed-Effects Models Using lme4. Journal of Statistical Software, 2015. 67: p. 1–48.

60. Lüdecke, D., et al., performance: An R Package for Assessment, Comparison and Testing of Statistical Models. Journal of Open Source Software, 2021. 6(60): p. 3139.

61. Lefcheck, J.S., piecewiseSEM: Piecewise structural equation modeling in R for ecology, evolution, and systematics. Methods in Ecology and Evolution, 2016. 7(5): p. 573–579.

62. Fernández-González, A.J., et al., Defining the root endosphere and rhizosphere microbiomes from the World Olive Germplasm Collection. Scientific Reports, 2019. 9: p. 20423.

63. Engblom, E., et al., Foam Spora in Running Waters of Southern Greenland. Polar Research, 1986. 4(1): p. 47–51.

64. Gönczöl, J. and Á. Révay, Fungal Spores in rainwater: stemflow, Throughfall and Gutter Conidial Assemblages. Fungal Diversity, 2004. 16: p. 67–86.

65. Orłowska, M., I. Lengiewicz, and M. Suszycka, Hyphomycetes Developing on Water Plants and Bulrushes in Fish Ponds. Polish Journal of Environmental Studies, 2004. 13(6): p. 703–707.

66. Al-Riyami, M., et al., Leaf Decomposition in a Mountain Stream in the Sultanate of Oman. International Review of Hydrobiology, 2009. 94(1): p. 16–28.

67. Kravetz, S., et al., The Genus Tetracladium in Pampean Streams (Buenos Aires, Argentina). Phytotaxa, 2018. 338(3): p. 276.

68. Hubbard, M., J.J. Germida, and V. Vujanovic, Fungal endophytes enhance wheat heat and drought tolerance in terms of grain yield and second-generation seed viability. Journal of Applied Microbiology, 2013. 116(1): p. 109–122.

69. Molina-Montenegro, M.A., et al., Antarctic root endophytes improve physiological performance and yield in crops under salt stress by enhanced energy production and Na+ sequestration. Scientific Reports, 2020. 10: p. 5819.

70. Pańka, D., et al., Production of phenolics and the emission of volatile organic compounds by perennial ryegrass (Lolium perenne L.)/Neotyphodium lolii association as a response to infection by Fusarium poae. Journal of Plant Physiology, 2013. 170(11): p. 1010–1019.

71. Xia, C., et al., Role of Epichloë Endophytes in Defense Responses of Cool-Season Grasses to Pathogens: A Review. Plant Disease, 2018. **102**(11).

72. Behie, S.W. and M.J. Bidochka, Nutrient transfer in plant–fungal symbioses. Trends in Plant Science, 2014. 19(11): p. 734–740.

73. Abrego, N., et al., Accounting for environmental variation in co-occurrence modelling reveals the importance of positive interactions in root-associated fungal communities. Molecular Ecology, 2020. 29(14): p. 2736–2746.

74. Kia, S.H., et al., Root endophytic fungi show low levels of interspecific competitioninplanta. Fungal Ecology, 2019. 39: p. 184–191.

75. Spagnoletti, F.N., et al., Dark septate endophytes present different potential to solubilize calcium, iron and aluminum phosphates. Applied Soil Ecology, 2017. 111: p. 25–32.

76. Sati, S.C. and P. Pant, Evaluation of Phosphate Solubilization by Root Endophytic Aquatic Hyphomycete Tetracladium setigerum. Symbiosis, 2018. 77(2): p. 141–145.

77. Wang, Y., et al., Different Selectivity in Fungal Communities Between Manure and Mineral Fertilizers: A Study in an Alkaline Soil After 30 Years Fertilization. Front. Microbiol., 2018. 9(2613).

78. Gosling, P., et al., Arbuscular mycorrhizal fungi and organic farming. Agriculture, Ecosystems & Environment, 2006. 113(1-4): p. 17–35.

79. Rousk, J., P.C. Brookes, and E. Bååth, Contrasting Soil pH Effects on Fungal and Bacterial Growth Suggest Functional Redundancy in Carbon Mineralization. Applied and Environmental Microbiology, 2009. 75(6): p. 1589–1696.

80. Grządziel, J. and A. Gałązka, Fungal Biodiversity of the Most Common Types of Polish Soil in a Long-Term Microplot Experiment. Front. Microbiol., 2019. 10(6).

81. Heintze, S.G., The use of the Glass Electrode in Soil Reaction and Oxidation-Reduction Potential Measurements. The Journal of Agricultural Science, 1934. 24(1): p. 28–41.

82. Husson, O., Redox potential (Eh) and pH as drivers of soil/plant/microorganism systems: a transdisciplinary overview pointing to integrative opportunities for agronomy. Plant Soil, 2012. 362: p. 389–417.

83. Włodarczyk, T., et al., Redox potential, nitrate content and pH in flooded Eutric Cambisol during nitrate reduction. Research in Agricultural Engineering 2007. 53(1).

84. Tano, B.F., et al., Spatial and Temporal Variability of Soil Redox Potential, pH and Electrical Conductivity across a Toposequence in the Savanna of West Africa. Agronomy Journal, 2020. 10.

85. Phillips, I.R., Phosphorus availability and sorption under alternating waterlogged and drying conditions. Communications in Soil Science and Plant Analysis 1998. 29(19-20).

86. Taheri, A.E., C. Hamel, and Y. Gan, Cropping practices impact fungal endophytes and pathogens in durum wheat roots. Applied Soil Ecology, 2016. 100: p. 104–111.

87. Benitez, M.-S., S.L. Osborne, and R.M. Lehman, Previous crop and rotation history effects on maize seedling health and associated rhizosphere microbiome. Scientific Reports, 2017. 15709.

88. Kirkegaard, J.A., et al., Effect of previous crops on crown rot and yield of durum and bread wheat in northern NSW. Australian Journal of Agricultural Research, 2004. 55(3): p. 321 – 334.

