## Supplementary figures and images for "Landscape scale ecology of *Tetracladium spp*. fungal root endophytes"

### Additional file 1

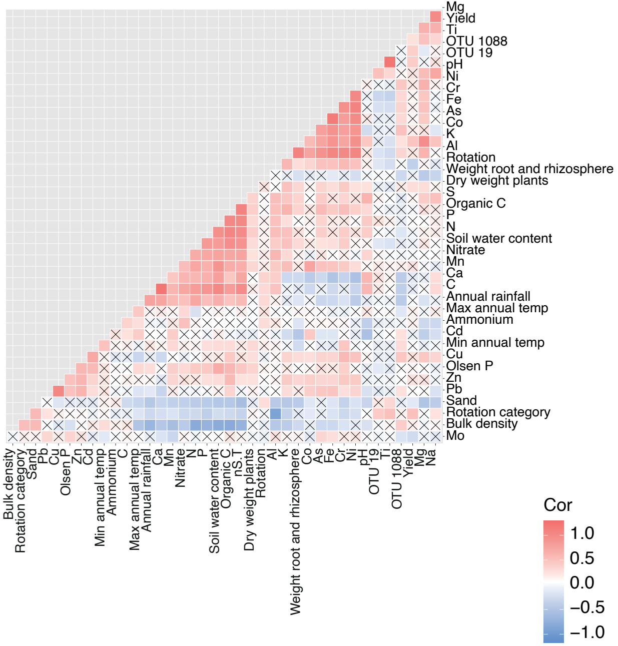
