## Additional file 2 for "Landscape scale ecology of *Tetracladium spp*. fungal root endophytes"

A

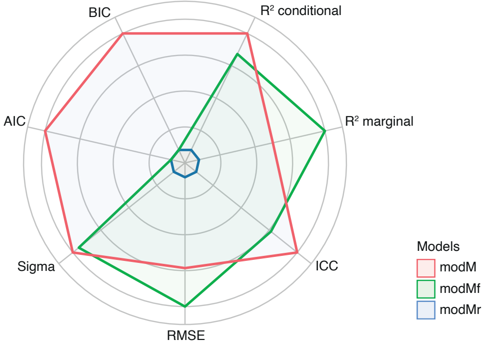


B

| Name | $R^2$ conditional | $R^2$ marginal | ICC | RMSE | Sigma | AIC | BIC | Performance score |
|---|---|---|---|---|---|---|---|---|
| modMr | 0.528 | 0.183 | 0.422 | 0.085 | 0.087 | <0.001 | <0.001 | 0.898 |
| modMf | 0.507 | 0.195 | 0.387 | 0.084 | 0.088 | <0.001 | <0.001 | 0.643 |
| modM | 0.409 | 0.167 | 0.290 | 0.088 | 0.090 | <0.001 | <0.001 | 0.000 |

C

| Name | $R^2$ conditional | $R^2$ marginal | ICC | RMSE | Sigma |
|---|---|---|---|---|---|
| modsoil | 0.413 | 0.011 | 0.407 | 0.089 | 0.091 |
| modclimate | 0.407 | 0.002 | 0.406 | 0.089 | 0.091 |
